# Network profiling of hepatocellular carcinoma targets for evidence based pharmacological approach to improve clinical efficacy

**DOI:** 10.1101/2022.02.21.481313

**Authors:** Bhavya Manchukonda, Arun HS Kumar

**Affiliations:** School of Medicine, Royal College of Surgeons in Ireland, Dublin, Ireland; Stemcology, School of Veterinary Medicine, University College Dublin, Belfield, Dublin-04, Ireland.

**Keywords:** Network pharmacology, Hepatocellular carcinoma, Clinical efficacy, network proteins, immunotherapy, Cell therapy

## Abstract

**Introduction:** Hepatocellular carcinoma (HCC) is the most prevalent malignancy of the liver with limited clinical efficacy of currently used drugs such as sorafenib. Hence in this study we assessed the network proteins of HCC targets to identify the target/s which can achieve optimal clinical efficacy.

**Materials and Methods:** The reported HCC targets and their network proteins were identified in the string database. The interactions of the network proteins based on the number of hydrogen bonds formed were evaluated using the chimera software and used to merit the network protein interactions. The merit of network protein interactions in clinical efficacy was assessed based on the expression pattern of the network proteins and corelating their targeting by sorafenib.

**Results:** 22 potential HCC targets were identified along with their 152 unique network proteins. The following HCC targets; PDGFRB, IFNA2, VEGFR2, PD1, C-MET, RAR and IGF1R were observed to be among the top networks with the most number of hydrogen bond interactions between them. Among these, C-MET, RAR and IGF1R were significantly expressed in hepatocytes, making them relevant HCC targets. PD-1 and PD-L1, which are immune checkpoint regulators and hence used as part of immune therapy, were observed to form higher numbers of hydrogen bonds with HCC network proteins.

**Conclusion:** Our analysis suggest that selectively targeting IGF1R, C-MET and RAR in hepatocytes together with immunotherapy will result in optimal clinical efficacy in the management of HCC.

## Introduction

Hepatocellular carcinoma (HCC) is the most prevalent malignancy of the liver and is often observed secondary to chronic liver conditions such as liver cirrhosis. In liver cirrhosis, healthy liver tissue is irreversibly replaced by fibrous scar tissue, impeding liver physiology^1, 2^. Several aetiologies which can lead to chronic liver pathologies also considerably increase the risk of developing HCC^2^. Screening for liver diseases in primary healthcare does not seem to be optimal as incidence of HCC has progressively increased in the last decade^3^. Early detection of HCC is key to disease management as it can be effectively treated by surgical approaches. Incidentally patients with HCC often present at later stages, when the disease has already advanced, limiting the merit of surgical approaches^4^. Moreover, it has been highlighted in a study that patients seem to have little to no knowledge regarding their liver disease prior to developing HCC^2^.

The survival rate at advanced stages of HCC is limited despite the availability of surgical and nonsurgical options for the disease management ^2^. The selection of these treatment options are based on the disease severity, metastasis and co-morbid clinical characteristics. Surgical options include surgical resection and liver transplant. The 5-year survival rates post-surgical resection are 40-70% with 70% recurrence rates. Whereas, liver transplant is often suggested for unresectable tumours which is often performed in combination with other nonsurgical treatments such as trans-arterial chemoembolization (TACE) or ablation. Nonsurgical treatment options are suggested when surgery is not suitable and these are divided into regional and systemic therapies^4^. Systemic therapies such as tyrosine kinase inhibitors (TKIs) are recommended in the management of advanced HCC cases. Sorafenib is one such TKI used as a first line of therapy. However, there are reports of tumour resistance development by various escape or compensation techniques, which reduce therapeutic efficacy. Additionally, immunotherapy agents such as bevacizumab are also being recommended, which help in strengthening innate immune response against tumour cells^4, 5^.

One of the major limitations with the currently available systemic therapies is the development of resistance or tolerance leading to relapse of the tumour^4^. Such therapeutic failures although less researched are often due to suboptimal targeting of the relevant tumour targets. Besides, the suboptimal targeting is directly a consequence of ignoring the influence of network proteins in tumour progression, as these network proteins play a vital role in therapeutic failure and tumour escape mechanisms^6, 7^. In our view a detailed study of the network proteins relevant to the progression of HCC will be helpful in identifying optimal therapeutic approaches to treating HCC. Hence in this study we focused on assessing the molecular interactions of HCC targets and their network proteins to suggest evidence based approaches to optimizing clinical efficacy of therapeutic targeting.

## Material and methods

The potential targets of HCC were identified from the Therapeutic Target Database (http://db.idrblab.net/ttd/). The network proteins of the HCC drug target proteins in humans were identified using the STRING database (https://string-db.org/). The PDB codes for all the proteins were selected through the UniProt database (https://www.uniprot.org/uniprot/P09619) using a specific selection criteria (i.e., based on the maximum length and resolution of the protein of interest). The PDB codes identified were cross verified from the protein data bank (https://www.rcsb.org/). The Chimera software was used to assess the hydrogen bond interactions between the HCC targets and each of the network proteins identified. From this analysis the protein interactions with the highest and the lowest number of hydrogen bonds were sub-selected for network profiling of HCC targets in this study. The interaction scores for the HCC target network proteins from the STRING database were documented. An analysis of the interaction score with that of the hydrogen bond between the network proteins was performed by sub-selecting the top 20 and lowest 20 interactive proteins from each of the approaches to interpret their biological relevance.

According to the American Society of Clinical Oncology (ASCO), the standard first-line of treatment protocol for HCC is atezolizumab and bevacizumab. Sorafenib or lenvatinib which are TKIs are recommended as first-line in cases where the use of atezolizumab and bevacizumab is contraindicated^8^. Consistent with this recommendation, the STITCH database (http://stitch.embl.de/) screening showed the following drugs; Cabozantinib S-malate, Lenvatinib Mesylate, Sorafenib Tosylate, Pemigatinib and Regorafenib within the primary network of HCC therapeutics. Hence the network proteins (Human specific) of these therapeutics along with their interaction scores were identified using the STITCH database. The HCC drugs were retrieved into the chimera software using their canonical smiles identity sequence from the PubChem database (https://pubchem.ncbi.nlm.nih.gov/). The expression profile of relevant HCC targets in human tissues was identified from the human protein atlas database (https://www.proteinatlas.org/about).

## Results

The Therapeutic Target Database search yielded 22 potential targets for HCC. The HCC targets are summarised in figure 1 along with a representative drug for each of the targets identified. The search for the networking proteins of these HCC targets showed 152 unique proteins in the STRING database with highly variable interaction scores (data not shown). Similarly the analysis of the hydrogen bond formation between the networking proteins also showed a considerable degree of variation (data not shown). Due to this inherent variability and vastness of the dataset, we assessed twenty protein networks with highest and lowest interaction scores or hydrogen bonds in this study.

**Figure 1:**
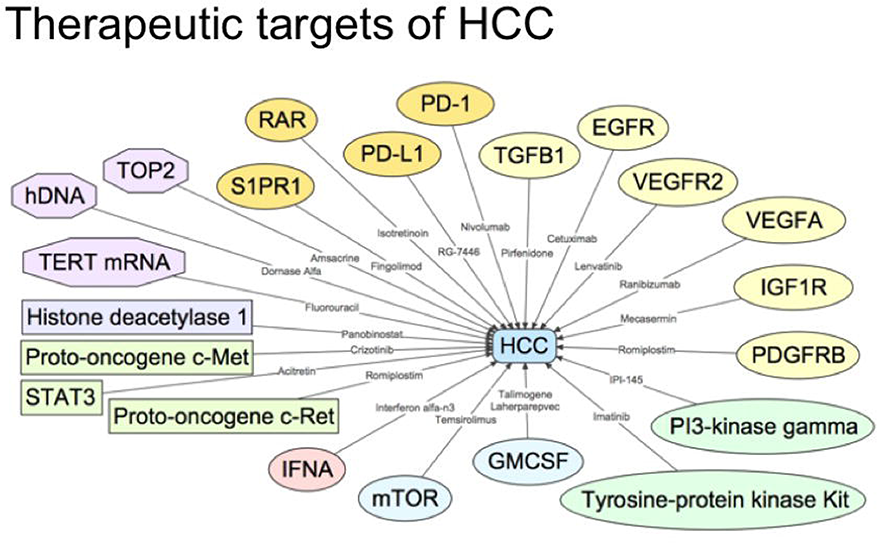
Summary of the therapeutic targets of hepatocellular carcinoma with a representative drug against each of the targets.

Among the interaction scores between protein networks, mTOR, VEGFA, VEGFR2, IGF1R, EGFR and HDAC1 were observed to be the top 20 networks with more than one protein interaction (Figure 2A). While the IFNA2, PD1, S1PR1 were prominent networks with least network interaction scores (Figure 2B). In contrast to the observations from the protein network interaction scores, the number of hydrogen bonds between protein networks were highest for the following HCC targets; PDGFRB, IFNA2, VEGFR2, PD1, P c-met, and IGF1R (Figure 2C). Additionally RAR-IRS2 network interaction showed over 20K hydrogen bonds, suggesting that these target interaction are biologically significant. It is interesting to note that VEGFR2 and IGF1R were the only two HCC targets correlating with both network interaction scores and the highest number of hydrogen bonds. The number of hydrogen bonds formed between the network proteins were least for the following HCC targets; P c-met, TGFb1 and RAR (Figure 2D).

**Figure 2:**
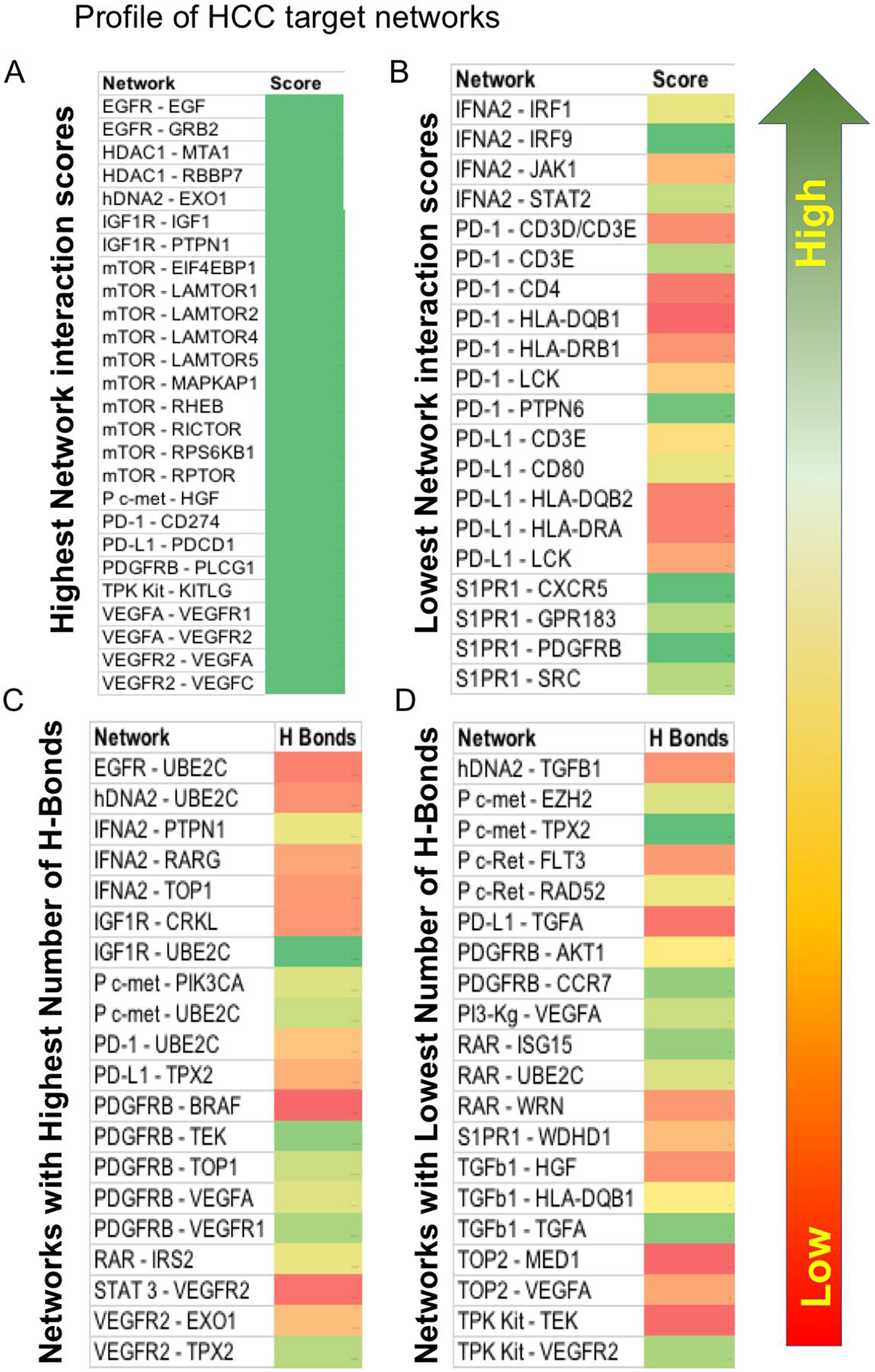
Profile of the hepatocellular carcinoma target networks. Network proteins with the highest (A) or lowest (B) network interactions scores in the String database. Network proteins with the highest (C) or lowest (D) number of hydrogen bonds between them. Arrow indicates the gradation of the colour code i.e., red and green indicating the lowest or highest number (interactions score or hydron bonds) respectively.

The HCC target protein PDGFRB had 5 major interactions among the top 20 network interactions. While the IFNA2 had 3 major interactions and IGF1R, VEGFR2 and P c-met had 2 major interactions. This suggests PDGFRB, IFNA2, IGF1R, VEGFR2 and P c-met play a major role in the pathogenesis of HCC compared to other network interactions observed. Also the failure to form biologically significant numbers of hydrogen bonds between the network proteins by mTOR, HDAC1, and VEGFA suggests that these HCC targets may not be biologically relevant in the pathogenesis of HCC.

The network pharmacology analysis of sorafenib, which is the first-line treatment for HCC, indicated that it targets only a minority of the top 20 network proteins (PDGFRB, KDR) relevant to HCC pathogenesis (Figure 3A). PDGFRB is one such protein that is targeted by sorafenib. However, in addition to PDGFRB, there are several other relevant network proteins observed among the top 20 list (IFNA2, VEGFR2, PD1, P c-met, IGF1R and RAR) which were not targeted by sorafenib. Moreover, the other proteins (KDR, FLT3, RET, KIT and RAF1) which are targeted by sorafenib were irrelevant to the pathogenesis of HCC, as these off-target proteins were not observed among the HCC targets/networks. This, perhaps, may explain the limited efficacy of sorafenib in the clinical management of HCC. Lenvatinib and cabozantinib, which are some of the other drugs used in the clinical management of HCC, showed suboptimal network interaction scores with PDGFRB in addition to influencing a few non-specific targets (Figure 3B).

**Figure 3:**
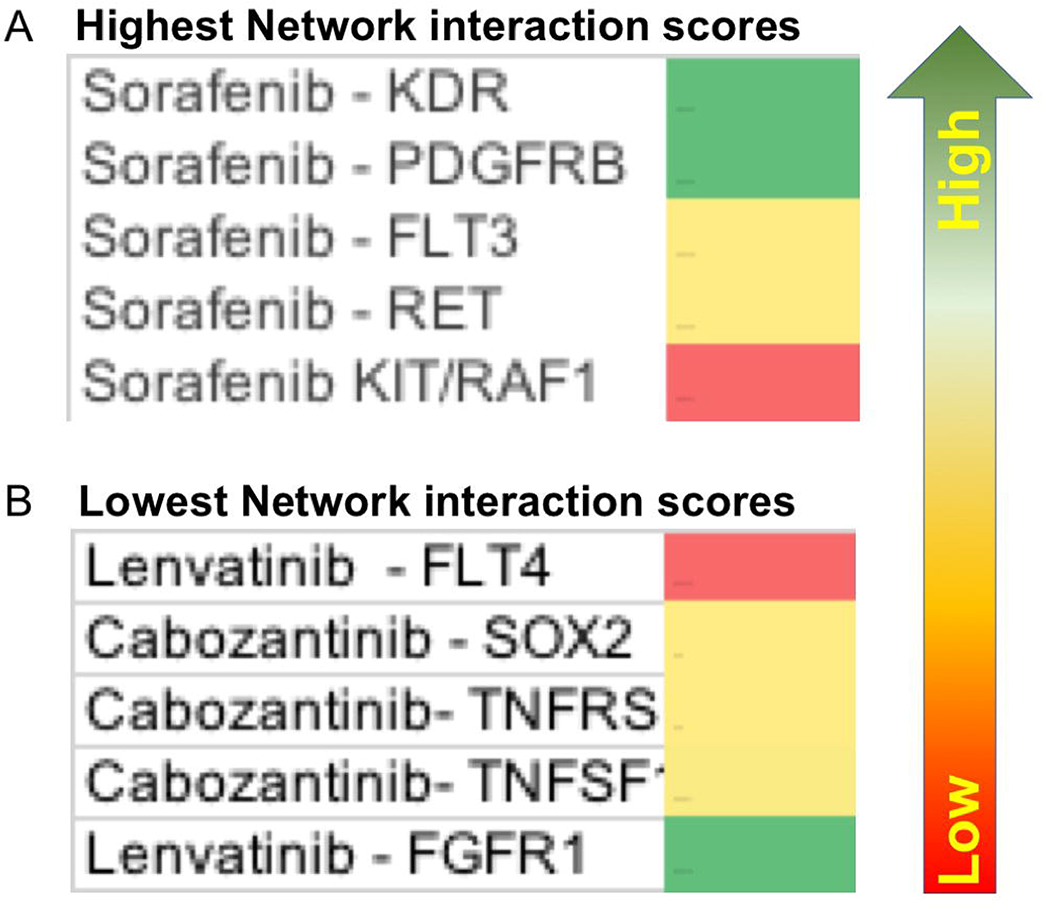
Network interactions scores of currently used drugs to treat hepatocellular carcinoma with their identified target. Arrow indicates the gradation of the colour code i.e., red and green indicating the lowest or highest interactions score respectively.

As the network analysis showed PDGFRB, IFNA2, VEGFR2, PD1, P c-met, and IGF1R to be relevant targets for HCC, we analysed their expression pattern in various human tissues. While PDGFRB, IFNA2, VEGFR2, PD1 were not expressed in hepatocytes. IGF1R (limited expression Intracellular and membrane), P c-met (vesicles and cytosol) and RAR (Nucleoplasm, nucleoli, actin filaments and cytosol) were expressed in hepatocytes to a variable degree (Figure 4).

**Figure 4:**
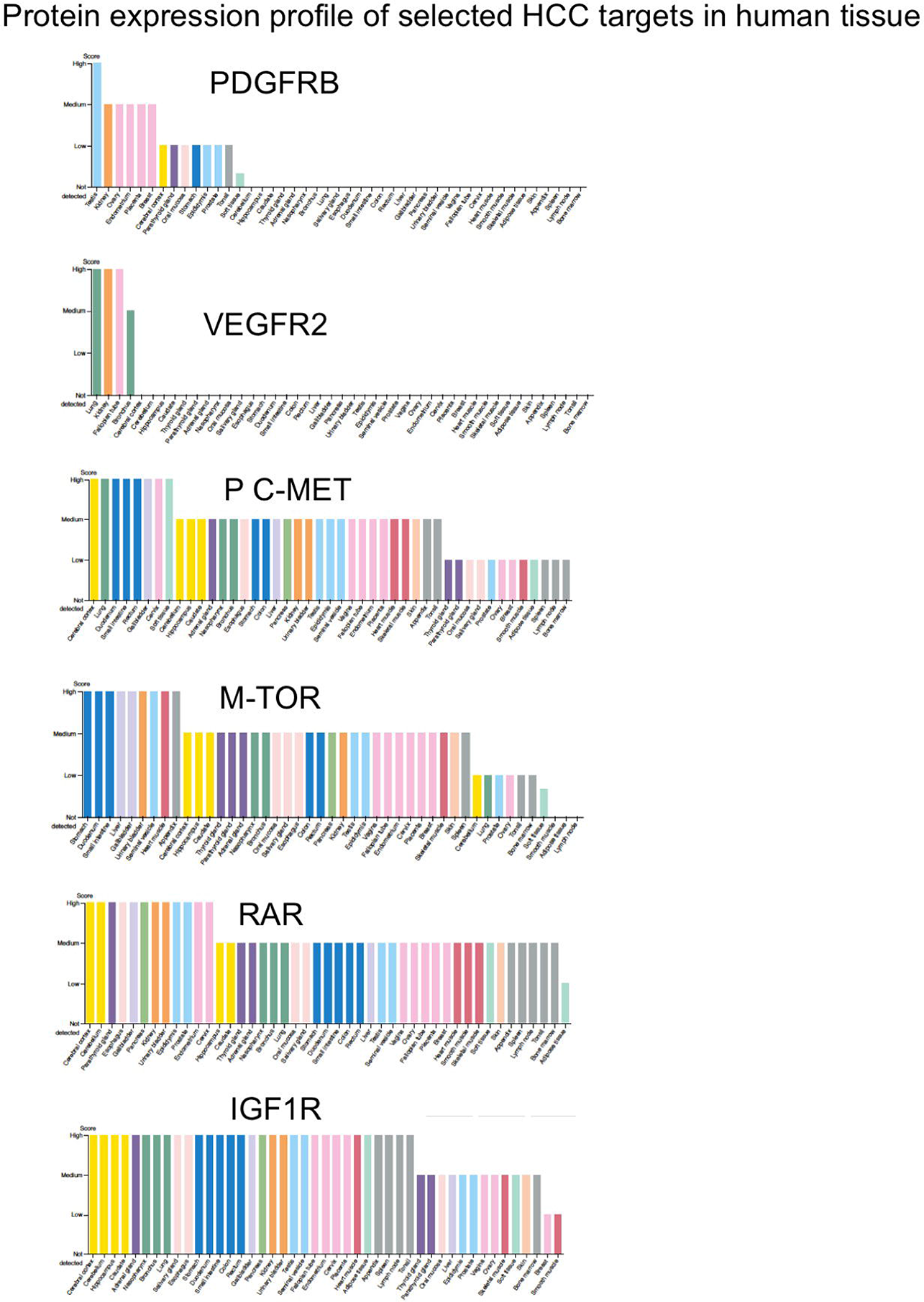
Expression profile of selected hepatocellular carcinoma targets in human tissues. Data sourced from the human protein atlas database (https://www.proteinatlas.org/about).

## Discussion

Proteins are highly social biomolecules which form biochemical networks to regulate various biological functions. Understanding of the protein networks relevant to specific disease conditions are valuable to designing optimal diagnostic and treatment strategies. Hence this study evaluated the protein networks of HCC targets with an aim to identify the optimal approaches to therapeutically target HCC progression. In this study PDGFRB, IFNA2, VEGFR2, PD1, P c-met, IGF1R and RAR were observed to be the most prominent HCC network protein based on the number of hydrogen bond interactions. Hydrogen bond interactions play a crucial role in the process of protein folding, structural confirmation and signalling which are necessary parts of establishing protein-protein interactions (PPI)^9,10^. Hence based on the significant hydrogen bond interactions between PDGFRB, IFNA2, VEGFR2, PD1, P c-met, IGF1R and RAR network proteins we can rationalise that these proteins play a vital role in progression of HCC.

Compared to the text mining approaches to study PPI, the approach of hydrogen bond interactions adopted in this study offers biologically relevant predictions on the influence of PPI on regulating physiology or pathology. Indeed in this study we observed that the PPI scores from STRING database analyses which are based on text mining approaches were ambiguous. and irrelevant to HCC targets as the highest 20 proteins (mTOR, VEGFA, VEGFR2, IGF1R, EGFR and HDAC1) identified in this study by this approach were not reported as therapeutically relevant targets for HCC. Hence we emphasise that validation of the PPI networks observed through text mining approaches by hydrogen bond interaction analysis is necessary to interpret biological relevance of PPI.

PDGFRB, IFNA2, VEGFR2, PD1, P c-met, IGF1R and RAR were the biological relevant protein networks identified in this study as essential to the progression of HCC. A prominent feature in the progression of HCC is the rapid and uncontrolled replication of hepatocytes with limited to negligible apoptosis processes. The network proteins identified in this study seem to pay a direct role in HCC progression by promoting hepatocyte replication (PDGFRB, VEGFR2, PD1, C-MET, RAR) while inhibiting apoptosis (IGF1R)^11–16^. PDGFRB is reported to regulate cell survival and migration, which are vital to progression of HCC^17^. Additionally PDGFRB also promoted angiogenesis which is essential to improve blood supply to the developing tumour^18, 19^. While these features of PDGFRB makes it an attractive target for HCC, but in the wider context of systemic physiology, targeting PDGFRB may result in several undesirable effects especially considering PDGFRB is poorly expressed in human hepatocytes. This may perhaps also explain poor efficacy of sorafenib, which primarily targets PDGFRB. Likewise VEGFR2 which plays a central role in angiogenesis, may also be of limited relevance as a therapeutic target for HCC.

IFNA2 is an interferon produced by macrophages potentially as a consequence to hepatitis caused by viruses. The antiviral, anti-inflammatory and antiproliferation activity IFNA2 can benefit in the context of HCC, where viral infections are the primary aetiology. However the major adverse effects associated with systemic administration of interferon is a concern which limits its therapeutic merit^20–23^. Nevertheless, with recent advancements in cell based therapies for cancers, interferon may have a role as priming agents in cell therapy protocols for optimising therapeutic efficacy, which warrants further research.

PD-1, commonly present on activated T-cells, binds to PD-L1 present on some cancer cells, which use this to escape destruction and promote the progression of cancer^24^. Our observation of higher numbers of hydrogen bonds between PD-1/UBE2C and PD-L1/TPX2 suggests that immune checkpoint inhibitors or immunotherapy will be beneficial to HCC patients. C-met was one of the most promising network proteins identified in our analysis. C-met together with IGF1R and RAR are the only network proteins expressed in hepatocytes in humans. C-met by signalling via the PI3K/AKT pathway promotes pro-survival of HCC. C-met is also reported to act as a receptor for entry of pathogens ^25, 26^, which supports its selective targeting in hepatocytes together with immune checkpoint inhibitors for optimal therapeutic efficacy in the management of HCC. In addition to supporting proliferation of HCC, IGF1R due to its anti-apoptotic effects further aids the survival of HCC, which is crucial to disease progression and severity. In this context we observed the highest number of hydrogen bonds between IGF1R and UBE2C, which further iterates the significance of targeting IGF1R for the clinical management of HCC.

RAR together with histone deacetylases tend to promote inflammatory response in endothelial cells^27, 28^. The relevance of RAR induced inflammatory process in the context of HCC progression is not known. In this study we observed a significant number of hydrogen bonds between RAR and IRS2, which suggests a potential role of RAR in HCC progression by influencing both proinflammatory and antiapoptotic pathways. Considering the higher expression of IGF1R, C-MET and RAR in hepatocytes and they being observed among the top 20 network proteins, we suggest that selectively targeting these three network proteins may help achieve optimal clinical efficacy in the management of HCC. Incidentally sorafenib which currently remains the first line of therapy for HCC doesn’t target any of these three relevant targets of HCC, which perhaps explains its limited clinical efficacy (i.e., under 30 percent of HCC patients). Further all the primary targets of sorafenib were not expressed in hepatocytes and with the exception of PDGFRB and VEGFR2 all other network proteins were observed to show the least number of hydrogen bond interactions between them. Besides, several patients are reported to develop resistance to sorafenib within 6 months of treatment ^29, 30^ and many suffer from several adverse effects such as hand-foot skin reactions, diarrhoea, hypertension and weight loss^31–33^. Considering the observations from this study, the suboptimal efficacy of sorafenib and its off target undesirable effects are not surprising.

The protein networks among the lowest 20 hydrogen bond interactions were TGFb, TPK Kit, PI3-kg, S1PR1, TOP2, and P c-ret and numerous drugs have been used in HCC treatment to target these proteins. However, based on our analysis, there is limited biological relevance in targeting these proteins as they have weak molecular interactions with network proteins relevant to HCC progression.

Despite the novel approach identified in this study to target HCC based on the interactions of network proteins, one of the limitations of this study is we didn’t look at all 152 network proteins and limited our analysis to the top 20 network proteins with most hydrogen bonds. Nevertheless the top 20 protein networks may be most relevant to progression of HCC especially when the relevant targets were highly expressed in hepatocytes. This suggests that our analysis has merit and selectively targeting IGF1R, C-MET and RAR in hepatocytes together with immunotherapy will result in optimal clinical efficacy in the management of HCC, which warrants validation by clinical trials.

## Acknowledgement

Research support from University College Dublin-Seed funding/Output Based Research Support Scheme (R19862, 2019), Royal Society-UK (IES\R2\181067, 2018) and Stemcology (STGY2708, 2020) is acknowledged.

